# A DNA barcode database for the woody plants of Japan

**DOI:** 10.1101/2021.11.11.468310

**Authors:** Suzuki Setsuko, Kensuke Yoshimura, Saneyoshi Ueno, James Raymond Peter Worth, Tokuko Ujino-Ihara, Toshio Katsuki, Shuichi Noshiro, Tomoyuki Fujii, Takahisa Arai, Hiroshi Yoshimaru

## Abstract

DNA barcode databases are increasingly available for a range of organisms facilitating the wide application of DNA barcode-based pursuits. Here we announce the development of a comprehensive DNA barcode database of the Japanese woody flora representing 43 orders, 99 families, 303 genera and 834 species and comprising 77.3% of genera and 72.2% of species of woody plants in Japan. A total of 6,216 plant specimens were collected from 223 sites (municipalities, i.e. city, town, village) across the subtropical, temperate, boreal and alpine biomes in Japan with most species represented by multiple accessions. This database utilised three chloroplast DNA regions (rbcL, trnH-psbA and matK) and consists of 14,404 barcode sequences. Individual regions varied in their identification rates with species-level and genus-level rates for rbcL, trnH-psbA and matK being 57.4%/ 96.2%, 78.5%/ 99.1 % and 67.8%/ 98%, respectively. Identification rates were higher using region combinations with total species level rates for two region combinations (rbcL & trnH, rbcL & matK, and trnH-psbA & matK) ranging between 90.6–95.8%, and for all three regions equal to 98.6%. Genus level identification rates were even higher ranging between 99.7–100% for two region combinations and being 100% for the three regions. These results indicate that this DNA barcode database is an effective resource for investigations of woody plants in Japan using DNA barcodes and provides a useful template for development of libraries for other components of the Japanese flora.

## 1 INTRODUCTION

DNA barcodes are short DNA fragments that are able to accurately and rapidly identify to the lowest taxonomic level possible (ideally to the species level) any unidentified organism including whole or fragmented specimens, wood, pollen, subfossils or environmental DNA. The ability to identify plant species via DNA barcoding has a great range of uses including for human health (such as identifying sources of pollen (Kraaijeveld *et al.*, 2015) or house dust (Craine *et al.*, 2017), in forensics (Ferri *et al.*, 2015), bio-security (Ashfaq & Hebert, 2016), nature conservation (such as environmental monitoring (Fahner *et al.*, 2016), biodiversity assessment (Burgess *et al.*, 2011) and enforcing trade laws of endangered species (Dormontt *et al.*, 2015)), agriculture (e.g. monitoring pollination and gene flow of crops (Richardson *et al.*, 2015) and various applications for scientific research (e.g. understanding past impacts of climate change (Giguet-Covex *et al.*, 2014) or for use in plant taxonomy and species discovery (Kress *et al.*, 2015)). Creating DNA barcode libraries, that is, a collection of DNA sequences associated with specimens that have verified taxonomic identifications (Kress *et al.*, 2015), is essential for use as a reference in order to identify unidentified samples. DNA barcoding libraries are now available for a wide range of organisms such as animals and fungi due to the availability of universal barcodes for these groups. However, unlike animals or fungi, there is no single universal DNA fragment for use in DNA barcoding of plants mostly due to the low level of mutation of organelle genomes in plants (Wolfe *et al.*, 1987). This has made it necessary to use multi-locus barcodes (Kress & Erickson, 2007) and in some cases to develop specific barcodes for the targeted plant species meaning that the development of DNA barcode libraries for plants is more complex and time consuming. Nonetheless, in the last decade such libraries have become available for plants of specific countries or regions (de Vere *et al.*, 2012; Kim *et al.*, 2012), specific taxonomic groups (Liu *et al.*, 2018; Nevill *et al.*, 2013) or individual biomes (Costion *et al.*, 2016; Saarela *et al.*, 2013).

Due to the enormous effort required to completely represent the full range of genetic diversity within species in DNA barcode libraries, especially for those covering many diverse taxa, the full range of sequence diversity may not be fully captured in DNA barcode libraries. This factor, together with low sequence divergence between closely related species, which is particularly common in species rich clades, along with taxonomic uncertainty, means that reliable identification to species level can be difficult (Parmentier *et al.*, 2013; Raupach *et al.*, 2014). However, for many applications of DNA barcoding assignment to higher taxonomic levels, such as the genus-level, is of considerable value and, in many plant groups, accuracy of assignment at this level is more reliable than at the species level (Wilson *et al.*, 2011).

In Japan, DNA barcode libraries have been developed for a range of taxonomic groups (Japanese DNA Barcode Database Committee, 2014). However, these have focussed exclusively on animals such as birds (Nishiumi, 2012), ticks (Takano *et al.*, 2014), snails (Hirose *et al.*, 2015) and snapping beetles (Oba *et al.*, 2015) with plants, excluding ferns (Ebihara *et al.*, 2010), so far having been overlooked. The Japanese archipelago has a highly diverse vascular plant flora with 6,000 species (of which approximately 1,000 are woody plants in ~100 families (Satake *et al.*, 1989)) of which around 2,900 are endemic (Biodiversity Center of Japan Nature Conservation Bureau Ministry of the Environment, 2010) and is one of the 35 hotspots of plant diversity in the world (Mittermeier *et al.*, 2004).

In this paper we announce the development of a DNA barcode database for nearly the entire woody plant flora of Japan (available on the publicly available Barcode of Life Data System (BOLD: https://www.boldsystems.org (Ratnasingham & Hebert, 2007)). This database consists of 6,216 specimens of woody plants (i.e. those plants with above ground perennial parts having lignified wood formed by secondary growth) sampled from 223 sites across the entire Japanese Archipelago representing subtropical, temperate, boreal and alpine biomes with multiple accessions collected across the range of each species where possible (Figure 1). We utilized three chloroplast regions (rbcL, matK and trnH-psbA) which have become widely used in plant DNA barcoding studies and have achieved high rates of species resolution (Burgess *et al.*, 2011; Kress *et al.*, 2009).

**Figure 1.**
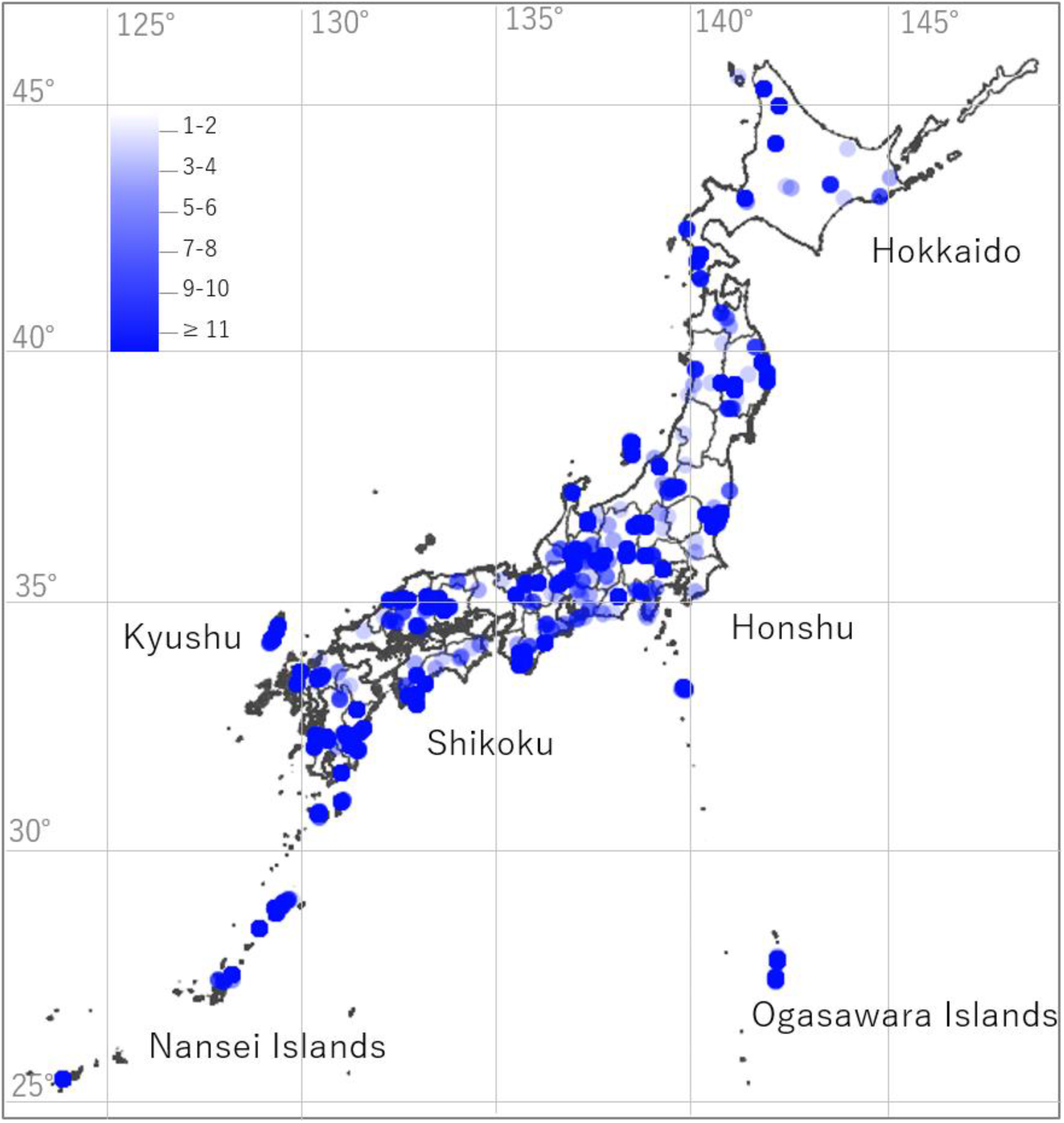
Location of all 223 sampling sites used to collect woody plants for the barcoding database. Opacity of the circles represent the number of samples per site ranging from 1 to 209 (average=24.9).

## 2 MATERIALS AND METHODS

### 2.1 Laboratory Work

Leaf samples for DNA extraction were collected from 223 sites across the whole of the Japanese Archipelago (Figure 1). These sites encompass all major vegetation types and biomes of Japan. For each sample, the latitude and longitude were recorded and identification was done to the lowest taxonomic level possible (i.e. subspecies or variety). For 92.4 % of samples, voucher herbarium specimens were prepared and stored in the herbaria in Japan. Most of them are housed in the xylarium (TWTw) and herbarium (TF) of the Forestry and Forest Products Research Institute (FFPRI), and most images of these voucher herbarium specimens are available at the database of Japanese Woods, FFPRI (https://db.ffpri.go.jp/WoodDB/index-E.html) and BOLD SYSTEMS. The rest vouchers of the samples are housed in the herbaria of Tohoku University Botanical Garden herbarium (TUS) and Herbarium of the Kyushu University Museum (FU) in Japan (Specimen details including museum ID together with GenBank ID and BOLD process ID is available from Supplementary table 1 and BOLD SYSTEMS).

**Table 1.**
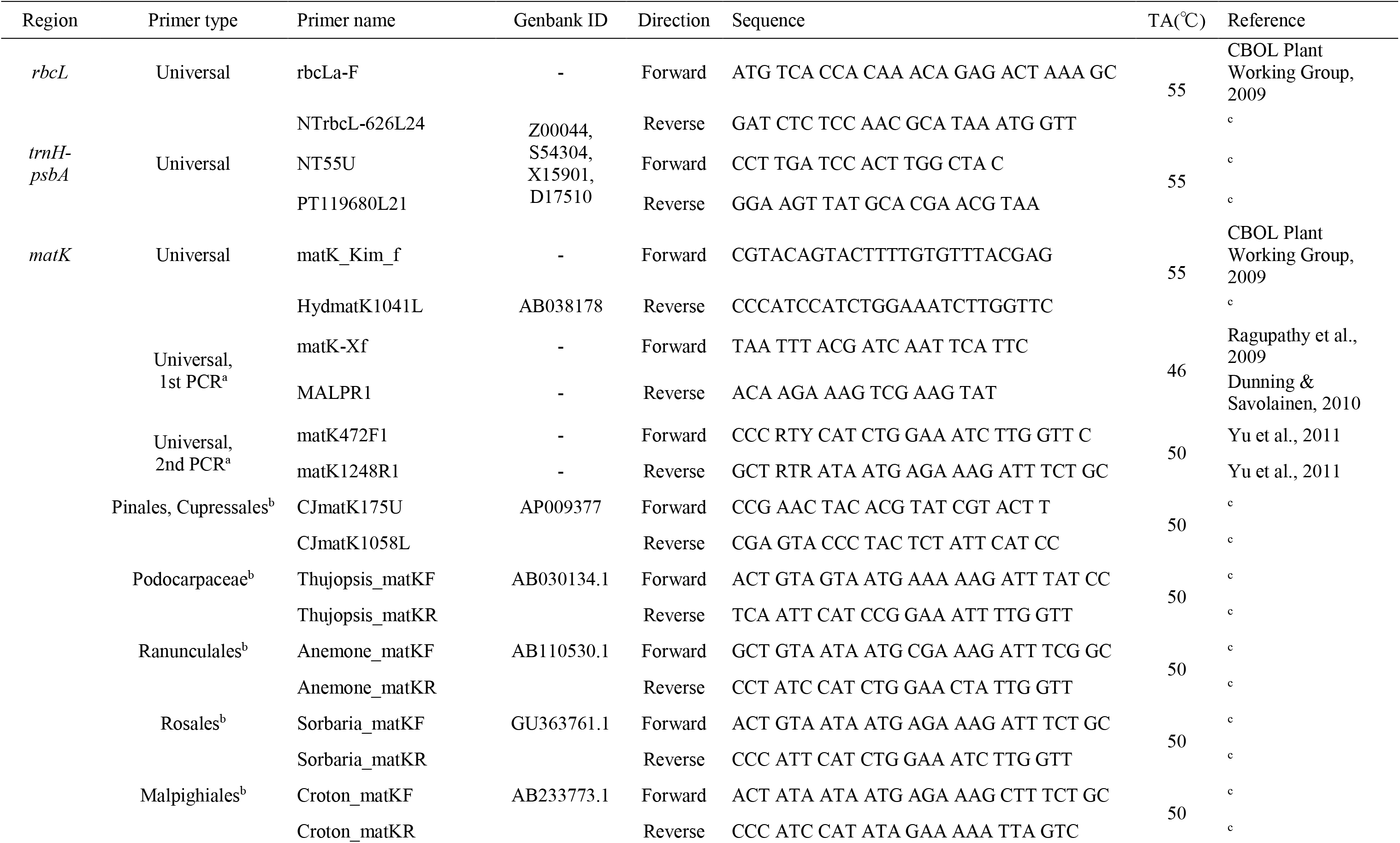
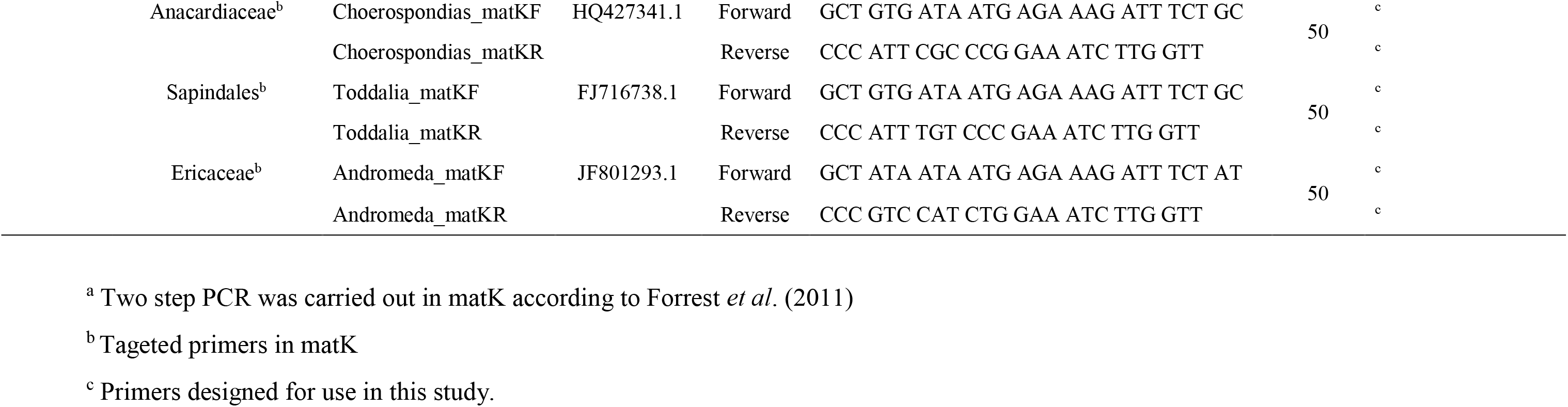
Details of the primer DNA sequences for each of the three chloroplast barcode regions.

DNA was extracted using DNeasy Plant Mini Kit (Qiagen) following the manufacturer’s instructions. For each sample we aimed to sequence three chloroplast barcode regions (rbcL, matK and trnH-psbA). The rbcL gene is easily amplified across land plants and, although it does not have sufficient variability to be used alone as a species discriminator, is considered a reliable ‘benchmark’ locus for placing taxa into family and genera (Kress & Erickson, 2007). matK is one of the most variable coding regions found in chloroplast DNA (Shaw *et al.*, 2005), while the non-coding trnH-psbA region has highly conserved priming sites across land plants and high sequence divergence (Kress & Erickson, 2007). The first two of these fragments, rbcL and matK, have been adopted by the Consortium for the Barcode of Life (CBOL) (CBOL Plant Working Group, 2009) as the core 2-locus barcode for plants while trnH-psbA has been widely used as a single locus or in combination with other loci (Pang *et al.*, 2012; Yao *et al.*, 2009). To amplify the barcode regions (rbcL, matK and trnH-psbA) we first tested primers recommended by the CBOL Plant Working Group (CBOL Plant Working Group, 2009; Hamilton, 1999). However, due to poor amplification and sequence quality, new primers to amplify both rbcL (reverse only) trnH-psbA (both forward and reverse) regions were designed by consulting the whole chloroplast genomes of tobacco, rice and Japanese black pine (Table 1). For the matK region, we also designed a new reverse primer by consulting the matK sequence of *Hydrangea macrophylla*, and trialled a two-step PCR approach following Forrest *et al*. (2011) with separate primer pairs for each step (Table 1). However, due to poor amplification success of matK in some orders or, in some cases, families, we developed new targeted primers. For those orders and/or families, where amplification of matK was poor we downloaded available chloroplast sequences from Genbank and developed new targeted primers (Table 1).

The PCR reaction mixture contained 0.05 μl Ex Taq polymerase (5 U/μl, TAKARA), 1μl 10X Ex Taq Buffer (20 mM, Mg^2+^ plus), 0.8 μl dNTP Mixture (2.5 mM each), 0.5 μl forward and reverse primer (each 2μM), and 2 μl template DNA (approximately 10 ng) in 10 μl total volume. PCRs were carried out using the following thermocycle: 94 °C for 3 min, then 35 cycles of 94 °C for 30s, each annealing temperature for 60s, 72 °C for 90s, followed by final extension at 72 °C for 10 min. Amplicon products were sent to TAKARA Bio (Mie, Japan) and Hokkaido System Science (Sapporo, Japan) for DNA sequencing or, alternatively, sequenced using an ABI3100 Genetic Analyzer (Applied Biosystems) at the FFPRI, Tsukuba, Ibaraki Prefecture. We used not only KB Basecaller (Applied Biosystems) but also PeakTrace software (Nucleics) for accurate base calling in some sequences. Sequences for rbcL and trnH were checked by eye using Sequencher (Hitachi) and aligned in Bioedit (Hall 1994). Those for matK were checked and aligned by CodonCode Aligner (CodonCode Corporation, www.codoncode.com).

### 2.2 Data analysis

Firstly, in order to grasp how well the database represents the total native woody plants of Japan we calculated the percentage of genera and species native to Japan included in the database per family using the most comprehensive reference available (The wild woody plants of Japan volumes I and II; Satake *et al.* 1989). In addition, we calculated the proportion of the flora included in the database for each of six regions of Japan (Hokkaido, Honshu, Shikoku, Kyushu, Nansei and Ogasawara Islands) with the distribution of each species based on Satake *et al.* (1989). Where new additions or changes to the woody plants have been made that are not listed in Satake *et al.* (1989) these were not taken into account in the calculations. For woody plants not listed in Satake *et al*. (1989), we included as many as possible from the 223 sites and included them in the database. For taxonomic classifications we followed the most recent available (Green List (Ito *et al.*, 2016) or, for those not listed on Green List, we followed YList (Yonekura & Kajita, 2003)). The taxonomic classification used for BOLD is shown in Supplementary table 1. Any disagreement in classification at the order or family level between Green List and/or YList is also indicated.

The success rate of sequencing was calculated as the proportion of the number of high-quality sequences obtained to the total number of samples. The species identification ability of each barcode was evaluated using the BLAST method (Altschul *et al.*, 1990). BLAST databases were constructed not only for each region (rbcL, matK and trnH-psbA), but also combined regions (rbcL & matK, matK & trnH-psbA, trnH-psbA & rbcL, and all three regions). Sequences were concatenated and used in the BLAST databases if sequences were available. Nucleotide BLAST (blastn) search was carried out for each sequence in each database against its own database (i.e. a self-blast) with default parameters. If the top hit sequence species name was the same as that of the query sequence and was the highest BLAST hit score, we considered the query sequence was successfully identified at the species level. However, if multiple top BLAST hits had the same score, the query was considered to not be identifiable to the species level. In the case that the multiple top BLAST hits contained only sequences of the same genus or family as that of query sequence, it was considered to be identified at genus or family level, respectively.

### 2.3 Phylogenetic analyses

To confirm the consistency of the data we constructed, a phylogenetic tree based on the three region barcode sequences of angiosperms included in the Japanese woody plant database and compared the result to published angiosperm phylogenies (The Angiosperm Phylogeny Group, 2003, 2009, 2016). To do so, for each angiosperm family one representative individual sample with the longest concatenated sequence of the three regions was selected. Sequence alignment was undertaken in Geneious version 2019.0.4 using Muscle alignment (Edgar, 2004) with default parameters. Phylogenetic analysis was undertaken using a Bayesian MCMC approach implemented in MrBayes v 3.2 (Ronquist *et al.*, 2012) and run with 400,000 MCMC generations, 4 chains and sample frequency of 100 and implementing the most parameter-rich substitution model, GTR+I+G, which has been shown to perform equally well as specifically selected models (Abadi *et al.*, 2019). A sequence of the basal angiosperm family Schisandraceae (Austrobaileyales) was selected as an outgroup (The Angiosperm Phylogeny Group, 2003, 2009, 2016). A consensus tree was produced using the sumt burnin=0.25 command and was then edited in FigTree v1.4.4 (Rambaut, 2020).

## 3 RESULTS

In total, 14,404 barcode sequences from 6,216 woody plant specimens were included in the database (Table 2). The average number of accessions per species was 7.5 and ranged from a single accession to a maximum of 73. Our database included 43 orders, 99 families, 303 genera, 834 species with 953 taxa which represented 77.3% of woody plant genera and 72.2% of woody plant species recognised by Satake *et al*. (1989). The missing species included those that are rare or have restricted ranges on islands, high mountain tops or serpentine regions or were otherwise not encountered at the 223 collection sites. The sampling rate for each geographic region ranged between 84.3–88.9% except for the Nansei Islands (63.3%, Table 3).

**Table 2.** A summary of the representation of genera and species of each plant order native to Japan included in the Japanese woody plants DNA barcoding database.

| Order | No. of genera in Japan<br>(% sampled)<sup>a</sup> |  | No. of species in Japan<br>(% sampled)<sup>a</sup> |  | No. taxa<br>sampled | No.<br>samples |
| --- | --- | --- | --- | --- | --- | --- |
| Cycadales | 1 | (100) | 1 | (100) | 1 | 2 |
| Pinales | 6 | (83) | 22 | (81.8) | 20 | 86 |
| Araucariales | 2 | (100) | 2 | (100) | 2 | 11 |
| Cupressales | 9 | (100) | 14 | (100) | 19 | 101 |
| Pandanales | 2 | (50) | 4 | (50) | 3 | 6 |
| Liliales<sup>b</sup> | - | - | - | - | 4 | 11 |
| Arecales | 6 | (50) | 6 | (50) | 4 | 10 |
| Poales | 6 | (0) | 15 | (0) | 0 | 0 |
| Ranunculales | 6 | (100) | 10 | (100) | 20 | 111 |
| Proteales | 3 | (66.7) | 7 | (71.4) | 5 | 42 |
| Trochodendrales | 1 | (100) | 1 | (100) | 1 | 10 |
| Buxales | 1 | (100) | 2 | (100) | 3 | 4 |
| Saxifragales | 9 | (88.9) | 24 | (66.7) | 20 | 115 |
| Vitales | 4 | (75) | 10 | (60) | 9 | 77 |
| Austrobaileyales | 3 | (100) | 4 | (100) | 5 | 43 |
| Fabales | 27 | (40.7) | 43 | (37.2) | 25 | 144 |
| Rosales | 42 | (88.1) | 172 | (73.8) | 155 | 1014 |
| Fagales | 14 | (100) | 56 | (94) | 60 | 555 |
| Cucurbitales | 1 | (100) | 1 | (100) | 1 | 6 |
| Celastrales | 5 | (100) | 27 | (74.1) | 23 | 169 |
| Oxalidales | 1 | (100) | 4 | (75) | 3 | 20 |
| Malpighiales | 28 | (60.7) | 68 | (60.3) | 43 | 215 |
| Myrtales | 11 | (63.6) | 16 | (56.25) | 10 | 23 |
| Crossosomatales | 4 | (100) | 4 | (100) | 8 | 93 |
| Chloranthales<sup>b</sup> | - | - | - | - | 1 | 3 |
| Sapindales | 20 | (90) | 61 | (86.9) | 68 | 629 |
| Malvales | 14 | (42.9) | 31 | (51.6) | 17 | 67 |
| Brassicales | 1 | (100) | 1 | (100) | 1 | 2 |
| Santalales | 7 | (85.7) | 9 | (77.8) | 7 | 37 |
| Caryophyllales | 1 | (100) | 3 | (33.3) | 1 | 2 |
| Cornales | 8 | (100) | 28 | (89.3) | 28 | 363 |
| Ericales | 31 | (90.3) | 159 | (74.2) | 146 | 844 |
| Icacinales | 2 | (0) | 2 | (0) | 0 | 0 |
| Garryales | 1 | (100) | 1 | (100) | 3 | 25 |
| Piperales<sup>b</sup> | - | - | - | - | 2 | 8 |
| Gentianales | 25 | (88) | 47 | (74.5) | 45 | 161 |
| Boraginales | 3 | (66.7) | 7 | (28.6) | 2 | 6 |
| Solanales | 3 | (33.3) | 6 | (16.7) | 1 | 2 |
| Lamiales | 15 | (80) | 49 | (75.5) | 45 | 265 |
| Aquifoliales | 2 | (100) | 24 | (91.7) | 28 | 171 |
| Asterales | 5 | (80) | 6 | (66.7) | 7 | 21 |
| Dipsacales | 7 | (100) | 51 | (68.6) | 45 | 274 |
| Apiales | 11 | (100) | 20 | (85) | 24 | 161 |
| Magnoliales | 2 | (50) | 7 | (85.7) | 8 | 54 |
| Laurales | 10 | (70) | 28 | (82.1) | 30 | 253 |
| Alismatales | 1 | (0) | 1 | (0) | 0 | 0 |
| Total | 361 | (77.3) | 1054 | (72.2) | 953 | 6216 |

<sup>a</sup> The percentage of genera and species native to Japan included in the database were calculated based on Satake *et al.* (1989).
<sup>b</sup> denotes families that are not included in Satake *et al.* (1989) and, therefore, the percentage of genera and species represented was not calculated. However, for these orders the number of taxa and species sampled are provided.

**Table 3.** Sampling rate for each region in Japan

| Region | No. of<br>species | No. of<br>species<br>sampled | Sampling<br>rate (%) |
| --- | --- | --- | --- |
| Hokkaido | 217 | 193 | 88.9 |
| Honshu | 664 | 560 | 84.3 |
| Shikoku | 495 | 441 | 89.1 |
| Kyushu | 527 | 454 | 86.1 |
| Nansei Islands | 420 | 266 | 63.3 |
| Ogsawara Islands | 88 | 76 | 86.4 |

The total sequence success for rbcL, trnH-psbA and matK were 96.2, 76.4 and 59.1%, respectively (Figure 2a). We obtained rbcL for all orders sampled while trnH-psbA did not amplify in one order (Araucariales, Figure 3a). The amplification of matK had been lower (45.0%) with no amplification in 14 orders consisting of 17 families including all gymnosperms when we used a two-step PCR approach (data not shown). However, the use of newly developed targeted primers (Table 1) resulted in the number of orders where matK was not amplifiable decreasing from 14 to 4 consisting of 10 families, and showed high sequence success rate in gymnosperms (Figure 3a, Supplementary table 2, 3).

**Figure 2.**
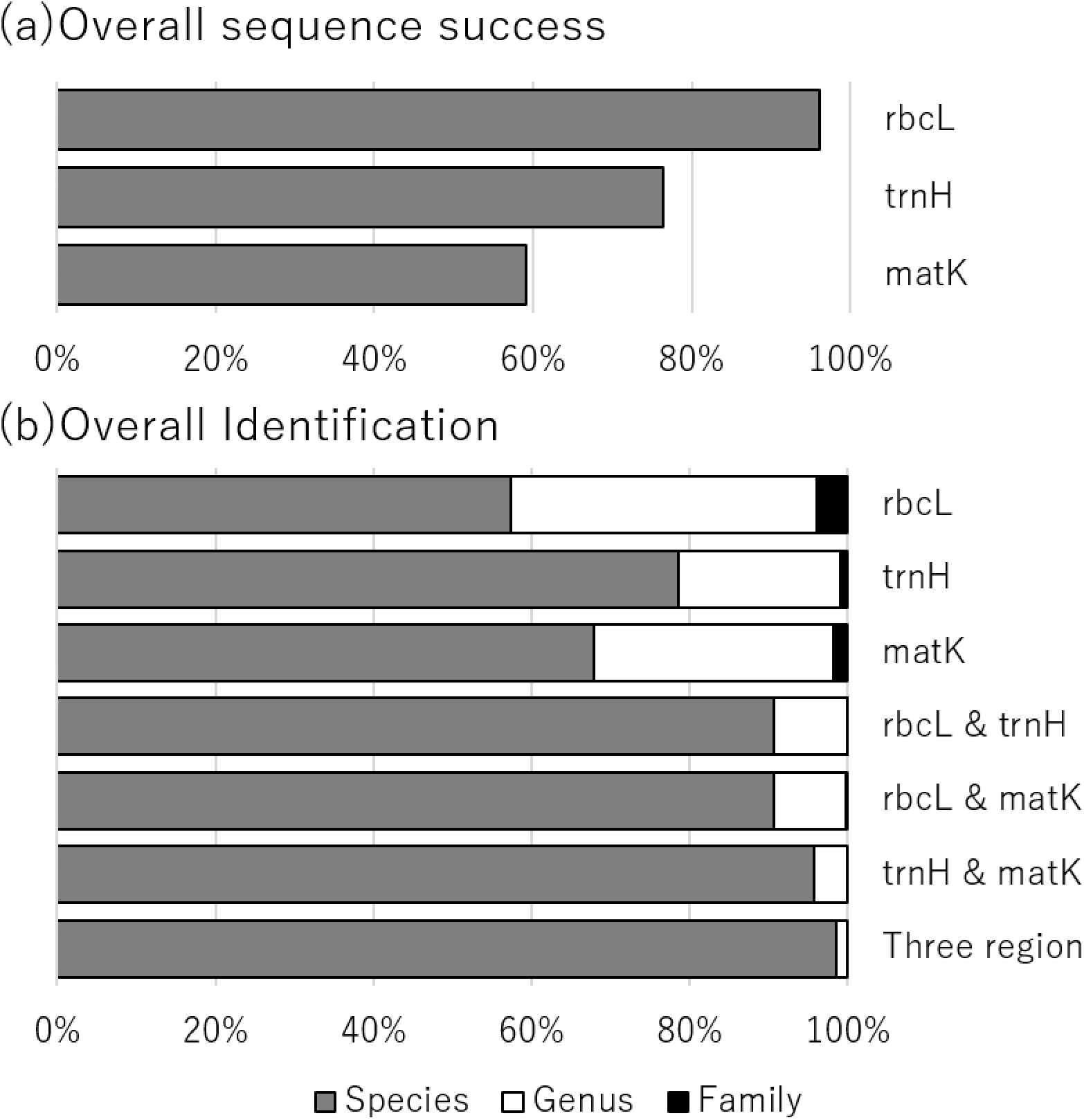
The overall level of sequencing success rate for each barcode region (a) and taxonomic identification rate (at the species, genus or family level) for each individual barcode region, barcode pair and all three regions (b).

**Figure 3.**
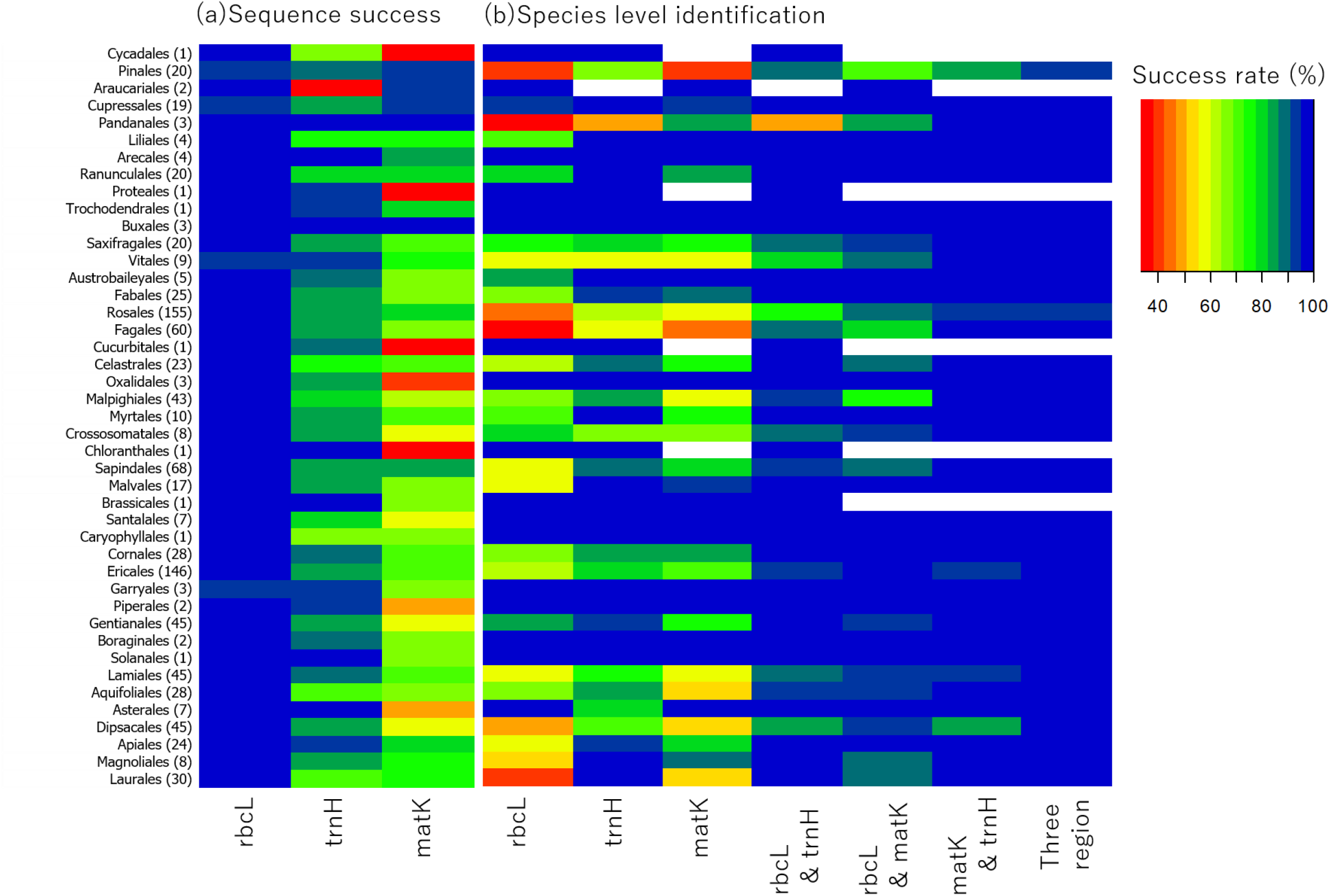
Sequencing success rate for each barcode region and species level identification rate for each barcode region, barcode pair and all three regions for each order. Number of taxa represented by an order is shown in brackets. White blanks show no sequence region available at the order.

Total identification rates for rbcL were as follows: 57.4% of samples returned a species level match, 96.2% a genus level match and 100% a family level match (Figure 2b). For trnH-psbA, there was greater identification rate with 78.5% returning a species level match, 99.1 % at genus level and 100% at the family level. For matK the identification rates were middle range with 67.8% returning a species level match, 98.1% at the genus level and 100% at the family level. Total identification rate at species level for two region combinations (rbcL & trnH, rbcL & matK, and trnH-psbA & matK) ranged between 90.6–95.8%, and for all three regions was 98.6%. Total identification rate at genus level for two region combinations (rbcL & trnH-psbA, rbcL & matK, and trnH-psbA & matK) ranged between 99.7–100%, and for three regions was 100 %.

There were some similar taxonomic patterns across the three regions for identification rate (Figure 3b). Some species rich orders had a tendency for low species level identification rate based on the individual regions such as Pinales, Pandanales, Rosales and Fagales. The lowest order-based identification rate for each region was 33.3% for rbcL, 50.0% for trnH (both Pandanales) and 40.5% for matK (Pinales) (Supplementary table 3). For 18 families that were well sampled and sequenced (i.e. over 70% of species represented and over 50% of sequence success for all three regions) and have relatively high species diversity (over 10), we found that identification rate at the species level was high using all three regions combined (Figure 4). Only two families had identification rate below 95% (Pinaceae=91.8% and Rosaceae=93.5%). Interestingly, despite low species-level identification rates based on each individual region (25.7–66.0%), the combined data resulted in a 99.2% successful identification rate for Fagaceae (Figure 4). High identification rates at each individual region of 85.4–97.8% were observed in Rutaceae. The identification rates at the family, genus and species levels for all families is provided in Supplementary table 3.

**Figure 4.**
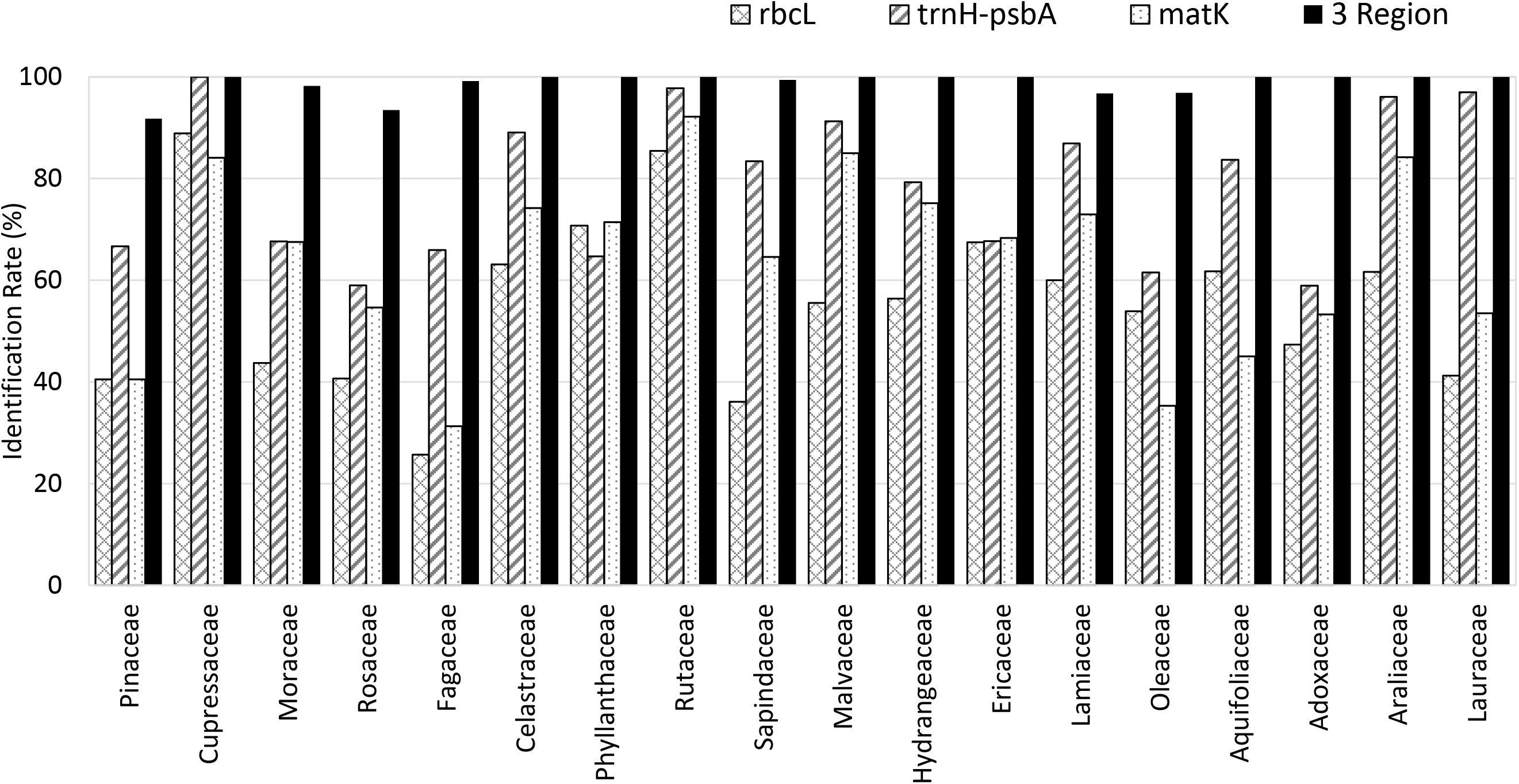
The species level identification rate for 18 selected plant families for which there are over 10 species in Japan and over 70% of species were sampled and over 50% sequence success for each three regions.

The phylogenetic tree of Japanese native angiosperm woody plants was well resolved with most nodes having branch-support values over 95%. In addition, the phylogenetic tree has similar overall relationships to published angiosperm phylogenies (Figure 5) (The Angiosperm Phylogeny Group, 2003, 2009, 2016).

**Figure 5.**
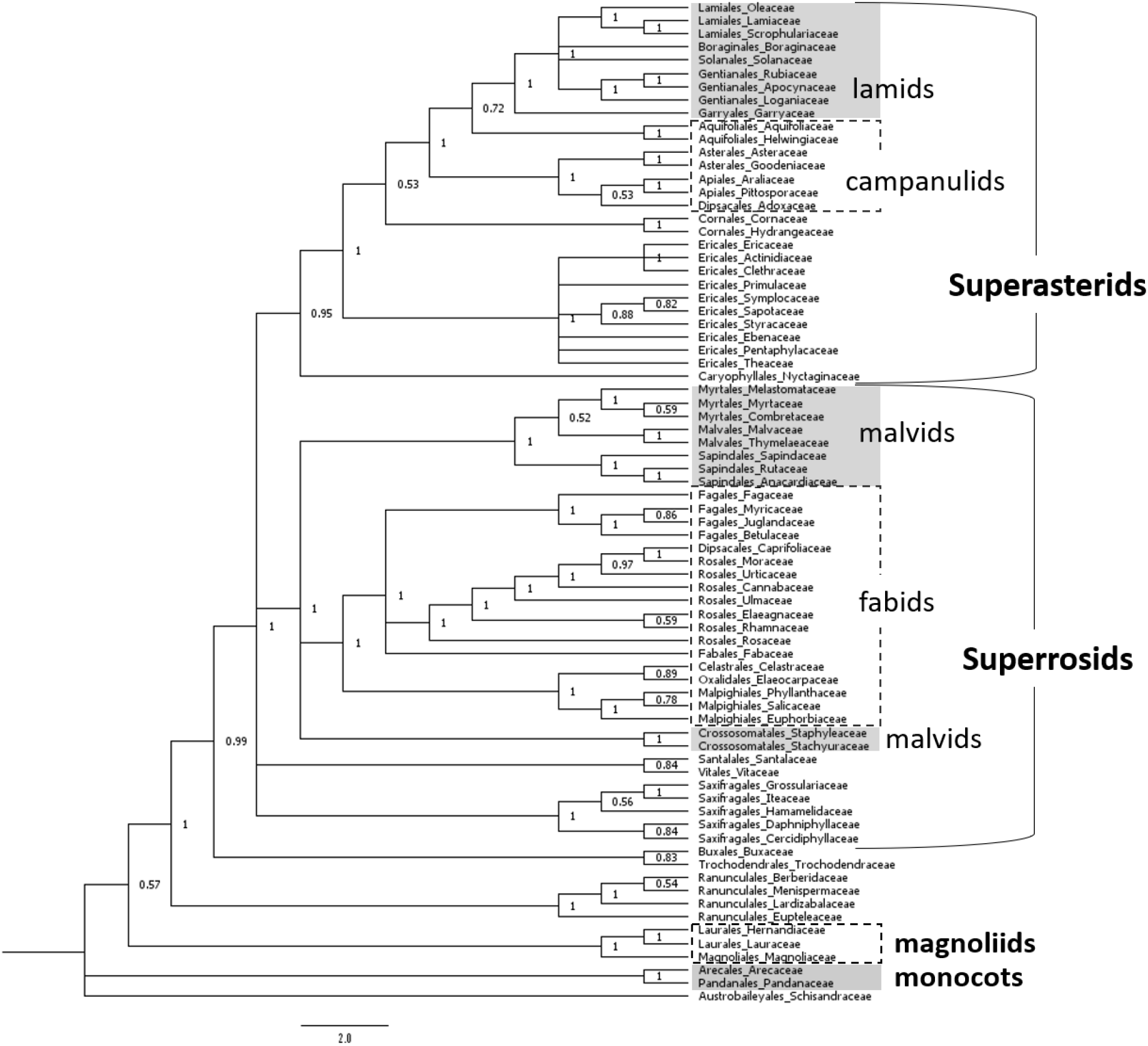
The phylogenetic tree of Japanese woody plants using all three barcode fragments (rbcL + trnH-psbA + matK).

## 4 DISCUSSION

This barcode database of Japanese woody plants provides a valuable resource for a range of applications in scientific, government and commercial pursuits. The high representation of woody plant species and multiple accessions per species make it one of the most comprehensive barcode libraries of any taxonomic group in the country to date. This database is significant advancement in terms of the barcode resources available for the Japanese flora with the only previous barcode database being for ferns (Ebihara *et al.*, 2010). For well sampled families the species identification rate achieved using all three barcodes was high (over 90%). The database can be accessed and samples analysed directly at the Barcode of Life Data System website (BOLD: Ratnasingham & Hebert (2007)), GenBank and ForestGEN (https://forestgen.ffpri.go.jp/jp/index.html) or alternatively, data exported and analysed in other programs (e.g. Sonet *et al*., 2013; Steinke *et al*., 2005; Vences *et al*., 2021). In the case that an unknown samples sequence is not included in the database identification to the nearest species and/or genus is likely to be accurate. Identification accuracy of unknown samples may be improved by utilising specific programs that take into account identification uncertainty due to incomplete sampling of sequence diversity (Sonet *et al.*, 2013).

Based on this database, the utility of each individual region differed substantially. rbcL had the highest sequence success rate but the lowest species-level identification rate. matK had the lowest sequence success rate and moderate species-level identification rate. In contrast, trnH-psbA had moderate sequencing success but the highest species-level identification rate. The CBOL Plant Working Group recommend the rbcL and matK combination as a standard barcode for land plants with trnH-psbA not included due to alignment issues caused by high variability including at mononucleotide repeats. However, if the application of barcodes is focussed at the family or genus level trnH-psbA is useful for species level identification given its high variability. On the hand, the rbcL and matK pair is more appropriate than trnH-psbA for revealing phylogenetic relationships across a diverse range of taxonomic groups. Lastly, we show that for orders where PCR amplification using universal primers was poor the development of targeted primers can significantly improve results. This was especially evident in gymnosperms and Sapindales where improvements of up to 100% were observed for matK (Supplementary table 2). This finding suggests that for reliable use of matK in DNA barcode libraries the development of targeted primers may be necessary.

The lowest species level identification rates based on individual regions were observed in Pinales, Pandanales, Rosales and Fagales. Rosales and Fagales have high species diversity but are also characterised by families where recurrent past or ongoing gene flow via hybridisation and/or hybrid species is widespread in Japan (Iwatsuki *et al.*, 1999, 2001, 2006). In contrast, Pinales have lower number of species but undergo extensive chloroplast sharing resulting in a lower identification rate in some genera (Aizawa & Iwaizumi, 2020; Watano *et al.*, 2004). Similar identification rates have been observed in European *Pinus* (Celiński *et al.*, 2017). Hybridisation between congeneric species can lead to chloroplast haplotype sharing (McKinnon *et al.*, 1999; Petit *et al.*, 2002) which in some cases makes species level identification difficult or even impossible, especially using a single chloroplast-based barcode. However, despite this even for orders with low species level identification rates based on individual regions such as Pinales, Rosales and Fagales, identification rates were over 90% using all three regions. Another reason for low species level identification rates observed in this study is exemplified by the genus *Pandanus* whose two Japanese species occur on separate islands but due to very recent speciation have not diverged at the chloroplast.

The availability of the Japanese woody plant barcode database is likely to open new avenues for scientific research and environmental monitoring in Japan. DNA Barcodes combined with NGS technology have already formed the basis of new, powerful and less invasive approaches to environmental monitoring in other organisms. For example, DNA barcodes for fish species used in metabarcoding of environmental DNA from seawater samples has the potential to revolutionize management of fish resources in Japan (Miya *et al.*, 2015) by improving the accuracy of assessments of fish species diversity (Yamamoto *et al.*, 2017) and providing a new tool to assess fish species biomass (Yamamoto *et al.*, 2016). Some research fields where DNA barcodes for Japanese woody plants may have significant impact, either as the sole investigative tool or together with existing methods, includes assessing present diversity of plants in biodiversity surveys (Taberlet *et al.*, 2012; Yoccoz *et al.*, 2012), diet analysis of endangered or invasive animals (Ando *et al.*, 2013; Nakahama *et al.*, 2021), or monitoring of pollen sources (Nakazawa *et al.*, 2013). The database could also have applications aimed at surveying past plant diversity from the decade to potentially thousands of years scale. For example, DNA metabarcoding could be a useful tool in deciphering the plant species composition of natural vegetation before conversion to plantations in the mid-late 20^th^ century. Reconversion to native forest and grasslands of some areas of under-exploited plantations in Japan has been an important issue in the last decade (Yamaura *et al.*, 2012) but given that plantation forests can cover large areas, with up to 66% of totals forest area in parts of southwestern Japan being planted forests (Forestry Agency, 2017), understanding past natural vegetation can be difficult. Chloroplast DNA barcodes are highly suited to investigations using degraded samples, including ancient DNA, because of the high copy number of chloroplast in plant cells (Wagner, 1992). However, given that shorter DNA fragments survive for longer (Deiner *et al.*, 2017), optimization of shorter targeted fragments of the three barcodes would most likely be required.

The barcode database for Japanese woody plants announced in this paper provides a valuable resource for both research and non-research-based pursuits investigating the countries flora. Due to the high species diversity and high number of geographically restricted species constructing a barcode database for the entire Japanese flora was not considered feasible in one study. However, we hope that this study provides a useful template from which further country-or region-based databases can be developed including for herbaceous plants and rare species that were not targeted in this study. For this database the use of nuclear loci such as the internal transcribed spacer (ITS) was not considered because of poor amplification success across land plants (Hollingsworth, 2011) and paralogous copies (Poczai & Hyvönen, 2010). However, future studies, especially those focussed on specific taxonomic groups, could have success using the shorter ITS2 region (China Plant B. O. L. Group *et al.*, 2011), or potentially other low copy nuclear loci (Kurian *et al.*, 2020).

## ACKNOWLEDGEMENTS

We would like to thank Y. Tsumura, T. Kawahara, M. Ohtani (FFPRI), M. Ito (The University of Tokyo) and H. Tachida (Kyushu University) for their valuable advice. We also thank, K. Ishida (Hirosaki University), H. Sakio, M. Nakata (Niigata University), N. Matsushita (The University of Tokyo), Y. Mukai, Shogo Kato (Gifu University), I. Tamaki, N. Yanagisawa (Gifu Academy of Forest Science and Culture), Y. Watanabe (Nagoya University), H. Kisanuki (Mie University), H. Ando, M. Yamazaki, S. Sakaguchi (Kyoto University), T. Yahara (Kyushu University), M. Takagi (University of Miyazaki), H. Taoda, K. Sugai, Shuri Kato, S. Kikuchi and T. Nagamitsu (FFPRI) for collecting samples and Y. Kawamata, A. Hisamatsu and C. Furusawa (FFPRI) for their assistance with laboratory work. This work was supported by a Grant-in-Aid for Scientific Research (20248017, 25292098) from the Japan Society for the Promotion of Science, and the support program of FFPRI for researchers having family responsibilities.

## AUTHOR CONTRIBUTIONS

Hiroshi Yoshimaru, Kensuke Yoshimura and Suzuki Setsuko conceived and designed the research. Toshio Katsuki, Shuichi Noshiro, Tomoyuki Fujii and Takahisa Arai conducted the field work. Hiroshi Yoshimaru, Kensuke Yoshimura, Takahisa Arai and Suzuki Setsuko conducted the laboratory work. Hiroshi Yoshimaru, Kensuke Yoshimura, Saneyoshi Ueno, James Raymond Peter Worth, Tokuko Ujino-Ihara and Suzuki Setsuko analyzed the data. James Raymond Peter Worth, Saneyoshi Ueno and Suzuki Setsuko wrote the manuscript. All authors contributed to the final submitted manuscript.

## DATA AVAILABILITY STATEMENT

Specimen data and DNA barcodes: BOLD and Genbank accessions are listed with specimen metadata in supporting information.

---

Supplementary table 1 List of samples used in this study

Can be downloaded from https://doi.org/10.6084/m9.figshare.16947391

Supplementary table 3 A summary of the representation of genera and species, sequence success of each barcode region, and identification rate at the speceis, genus and family level for each plant family native to Japan included in the Japanese Woody Plant DNA Barcoding Database.

Can be downloaded from https://doi.org/10.6084/m9.figshare.16947391

**Supplementary table 2.** Sequencing success rates of matK using two-step PCR and order specific primers

| Order | Family | Primer name | Sequence success rate for two-step PCR (%) | Sequence success rate for order specific primers (%) |
| --- | --- | --- | --- | --- |
| Pinales | Pinaceae | CJmatK | 0.00 | 89.6 |
| Araucariales | Podocarpaceae | Thujopsis_matK | 0.00 | 93.3 |
| Cupressales | Cupressaceae | CJmatK | 0.00 | 94.9 |
|  | Sciadopityaceae | CJmatK | 0.00 | 100.0 |
|  | Taxaceae | CJmatK | 0.00 | 92.9 |
| Ranunculales | Berberidaceae | Anemone_matK | 44.44 | 52.6 |
|  | Lardizabalaceae | Anemone_matK | 64.71 | 100.0 |
|  | Menispermaceae | Anemone_matK | 69.57 | 19.0 |
|  | Ranunculaceae | Anemone_matK | 53.33 | 96.6 |
| Rosales | Rosaceae | Sorbaria_matK | 34.33 | 93.1 |
| Malpighiales | Euphorbiaceae | Croton_matK | 14.63 | 40.8 |
| Sapindales | Anacardiaceae | Choerospondias_matK | 42.86 | 93.0 |
|  | Meliaceae | Toddalia_matK | 0.00 | 100.0 |
|  | Rutaceae | Toddalia_matK | 17.48 | 94.8 |
|  | Simaroubaceae | Toddalia_matK | 0.00 | 100.0 |
| Ericales | Ericaceae | Andromeda_matK | 37.54 | 87.7 |

## REFERENCES

Abadi S, Azouri D, Pupko T, Mayrose I (2019) Model selection may not be a mandatory step for phylogeny reconstruction. Nature communications, 10(1), 934.

Aizawa M, Iwaizumi MG (2020) Natural hybridization and introgression of *Abies firma* and *Abies homolepis* along the altitudinal gradient and genetic insights into the origin of *Abies umbellata*. Plant Species Biology, 35(2), 147–157.

Altschul SF, Gish W, Miller W, Myers EW, Lipman DJ (1990) Basic local alignment search tool. J Mol Biol, 215(3), 403–410.

Ando H, Setsuko S, Horikoshi K, et al. (2013) Diet analysis by next-generation sequencing indicates the frequent consumption of introduced plants by the critically endangered red-headed wood pigeon (*Columba janthina nitens*) in oceanic island habitats. Ecol Evol, 3(12), 4057–4069.

Ashfaq M, Hebert PDN (2016) DNA barcodes for bio-surveillance: regulated and economically important arthropod plant pests. Genome, 59(11), 933–945.

Biodiversity Center of Japan Nature Conservation Bureau Ministry of the Environment (2010) Biodiversity of Japan : a harmonious coexistence between nature and humankind. Tokyo: Heibonsha.

Burgess KS, Fazekas AJ, Kesanakurti PR, et al. (2011) Discriminating plant species in a local temperate flora using the rbcL+ matK DNA barcode. Methods in Ecology and Evolution, 2(4), 333–340.

CBOL Plant Working Group (2009) A DNA barcode for land plants. Proc Natl Acad Sci U S A, 106(31), 12794–12797.

Celiński K, Kijak H, Wojnicka-Półtorak A, et al. (2017) Effectiveness of the DNA barcoding approach for closely related conifers discrimination: A case study of the *Pinus mugo* complex. Comptes Rendus Biologies, 340(6), 339–348.

China Plant B. O. L. Group, Li D-Z, Gao L-M, et al. (2011) Comparative analysis of a large dataset indicates that internal transcribed spacer (ITS) should be incorporated into the core barcode for seed plants. Proceedings of the National Academy of Sciences, 108(49), 19641–19646.

Costion CM, Lowe AJ, Rossetto M, et al. (2016) Building a plant DNA barcode reference library for a diverse tropical Flora: an example from Queensland, Australia. Diversity, 8(1), 5–5.

Craine JM, Barberán A, Lynch RC, et al. (2017) Molecular analysis of environmental plant DNA in house dust across the United States. Aerobiologia, 33(1), 71–86.

de Vere N, Rich TCG, Ford CR, et al. (2012) DNA barcoding the native flowering plants and conifers of Wales. PLoS One, 7(6).

Deiner K, Bik HM, Mächler E, et al. (2017) Environmental DNA metabarcoding: Transforming how we survey animal and plant communities. Molecular ecology, 26(21), 5872–5895.

Dormontt EE, Boner M, Braun B, et al. (2015) Forensic timber identification: It’s time to integrate disciplines to combat illegal logging. Biological Conservation, 191, 790–798.

Dunning LT, Savolainen V. (2010) Broad-scale amplification of matK for DNA barcoding plants, a technical note. Botanical Journal of the Linnean Society, 164(1), 1–9.

Ebihara A, Nitta JH, Ito M (2010) Molecular species identification with rich floristic sampling: DNA barcoding the pteridophyte flora of Japan. PLoS One, 5(12), e15136.

Edgar RC (2004) MUSCLE: multiple sequence alignment with high accuracy and high throughput. Nucleic Acids Research, 32(5), 1792–1797.

Fahner NA, Shokralla S, Baird DJ, Hajibabaei M (2016) Large-scale monitoring of plants through environmental DNA metabarcoding of soil: recovery, resolution, and annotation of four DNA markers. PLoS One, 11(6).

Ferri G, Corradini B, Ferrari F, et al. (2015) Forensic botany II, DNA barcode for land plants: Which markers after the international agreement? Forensic Science International: Genetics, 15, 131–136.

Forestry Agency (2017) Status of forest resources. Proportion of forest and planted forest for each prefectures in Japan. Retrieved from https://www.rinya.maff.go.jp/j/keikaku/genkyou/h29/1.html

Forrest A, Hollingsworth P, Little D, et al. (2011) Plant DNA Barcoding using matK, some work in new primer sets Retrieved from https://www.slideshare.net/CBOLAdelaide2011/thursday-napier-lg29-1100-hollingsworth-mat-k-primers.

Giguet-Covex C, Pansu J, Arnaud F, et al. (2014) Long livestock farming history and human landscape shaping revealed by lake sediment DNA. Nature communications, 5(1), 1–7.

Hamilton MB (1999) Four primer pairs for the amplification of chloroplast intergenic regions with intraspecific variation. Molecular Ecology, 8(3), 521–523.

Hirose M, Hirose E, Kiyomoto M (2015) Identification of five species of *Dendrodoris* (Mollusca: Nudibranchia) from Japan, using DNA barcode and larval characters. Marine Biodiversity, 45(4), 769–780.

Hollingsworth PM (2011) Refining the DNA barcode for land plants. Proceedings of the National Academy of Sciences, 108(49), 19451–19452.

Ito M, Nagamasu H, Fujii S, et al. (2016) GreenList ver. 1.01. Retrieved from http://www.rdplants.org/gl/

Iwatsuki K, Boufford D, Ohba H (1999) Flora of Japan Vol. IIc. Angiospermae Dicotyledoneae Archlamydeae (c).

Iwatsuki K, Boufford D, Ohba H (2001) Flora of Japan Vol. IIb. Angiospermae Dicotyledoneae Archlamydeae (b). In: Kodansha, Tokyo.

Iwatsuki K, Boufford D, Ohba H (2006) Flora of Japan Vol. IIa. Angiospermae Dicotyledoneae Archlamydeae (a). In: Kodansha, Tokyo.

Japanese DNA Barcode Database Committee (2014) Japanese DNA Barcode Database (JBOL-DB). Retrieved from http://db.jboli.org/?locale=en

Kim S, Kim C-B, Min G-S, et al. (2012) Korea barcode of life database system (KBOL). Animal cells and systems, 16(1), 11–19.

Kraaijeveld K, De Weger LA, Ventayol García M, et al. (2015) Efficient and sensitive identification and quantification of airborne pollen using next‐generation DNA sequencing. Molecular Ecology Resources, 15(1), 8–16.

Kress WJ, Erickson DL (2007) A two-locus global DNA barcode for land plants: the coding rbcL gene complements the non-coding trnH-psbA spacer region. PLoS One, 2(6).

Kress WJ, Erickson DL, Jones FA, et al. (2009) Plant DNA barcodes and a community phylogeny of a tropical forest dynamics plot in Panama. Proceedings of the National Academy of Sciences, 106(44), 18621–18626.

Kress WJ, García-Robledo C, Uriarte M, Erickson DL (2015) DNA barcodes for ecology, evolution, and conservation. Trends in ecology & evolution, 30(1), 25–35.

Kurian A, Dev SA, Sreekumar VB, Muralidharan EM (2020) The low copy nuclear region, RPB2 as a novel DNA barcode region for species identification in the rattan genus *Calamus* (Arecaceae). Physiology and molecular biology of plants : an international journal of functional plant biology, 26(9), 1875–1887.

Liu J, Milne RI, Möller M, et al. (2018) Integrating a comprehensive DNA barcode reference library with a global map of yews (Taxus L.) for forensic identification. Molecular Ecology Resources, 18(5), 1115–1131.

McKinnon GE, Steane DA, Potts BM, Vaillancourt RE (1999) Incongruence between chloroplast and species phylogenies in Eucalyptus subgenus Monocalyptus (Myrtaceae). American Journal of Botany, 86(7), 1038–1046.

Mittermeier RA, Gil PR, Hoffman M, et al. (2004) Hotspots Revisited: Earth’s Biologically Richest and Most Endangered Terrestrial Ecoregions Cemex.

Miya M, Sato Y, Fukunaga T, et al. (2015) MiFish, a set of universal PCR primers for metabarcoding environmental DNA from fishes: detection of more than 230 subtropical marine species. Royal Society Open Science, 2(7), 150088.

Nakahama N, Furuta T, Ando H, et al. (2021) DNA meta-barcoding revealed that sika deer foraging strategies vary with season in a forest with degraded understory vegetation. Forest Ecology and Management, 484, 118637.

Nakazawa F, Uetake J, Suyama Y, et al. (2013) DNA analysis for section identification of individual Pinus pollen grains from Belukha glacier, Altai Mountains, Russia. Environmental Research Letters, 8, 014032.

Nevill PG, Wallace MJ, Miller JT, Krauss SL (2013) DNA barcoding for conservation, seed banking and ecological restoration of Acacia in the Midwest of Western Australia. Molecular Ecology Resources, 13(6), 1033–1042.

Nishiumi I (2012) DNA barcoding and species classification of Japanese birds. Japanese Journal of Ornithology, 61(2), 223–237.

Oba Y, Ôhira H, Murase Y, Moriyama A, Kumazawa Y (2015) DNA barcoding of Japanese click beetles (Coleoptera, Elateridae). PLoS One, 10(1), e0116612.

Pang X, Liu C, Shi L, et al. (2012) Utility of the trnH–psbA intergenic spacer region and its combinations as plant DNA barcodes: a meta-analysis. PLoS One, 7(11).

Parmentier I, Duminil Jm, Kuzmina M, et al. (2013) How effective are DNA barcodes in the identification of African rainforest trees? PLoS One, 8(4).

Petit RJ, Csaikl UM, Bordács S, et al. (2002) Chloroplast DNA variation in European white oaks: phylogeography and patterns of diversity based on data from over 2600 populations. Forest Ecology and Management, 156(1–3), 5–26.

Poczai P, Hyvönen J (2010) Nuclear ribosomal spacer regions in plant phylogenetics: problems and prospects. Molecular biology reports, 37(4), 1897–1912.

Ragupathy S, Newmaster SG, Murugesan M, Balasubramaniam V (2009) DNA barcoding discriminates a new cryptic grass species revealed in an ethnobotany study by the hill tribes of the Western Ghats in southern India. Molecular Ecology Resources, 9, 164–171.

Rambaut A (2020) FigTree v1.4.4. Institute of Evolutionary Biology, University of Edinburgh, Edinburgh. Retrieved from http://tree.bio.ed.ac.uk/software/figtree/

Ratnasingham S, Hebert PDN (2007) BOLD: The Barcode of Life Data System (http://www.barcodinglife.org). Molecular Ecology Notes, 7(3), 355–364.

Raupach MJ, Hendrich L, Küchler SM, et al. (2014) Building-up of a DNA barcode library for true bugs (Insecta: Hemiptera: Heteroptera) of Germany reveals taxonomic uncertainties and surprises. PLoS One, 9(9).

Richardson RT, Lin CH, Sponsler DB, et al. (2015) Application of ITS2 metabarcoding to determine the provenance of pollen collected by honey bees in an agroecosystem. Applications in plant sciences, 3(1), 1400066–1400066.

Ronquist F, Teslenko M, van der Mark P, et al. (2012) MrBayes 3.2: efficient Bayesian phylogenetic inference and model choice across a large model space. Syst Biol, 61(3), 539–542.

Saarela JM, Sokoloff PC, Gillespie LJ, Consaul LL, Bull RD (2013) DNA barcoding the Canadian Arctic flora: core plastid barcodes (rbcL+ matK) for 490 vascular plant species. PLoS One, 8(10).

Satake Y, Hara H, Watari S, Tominari T (1989) Wild flowers of Japan: Woody plants I, II. Tokyo: Heibonsha Ltd., Publishers. In: Japanese.

Shaw J, Lickey EB, Beck JT, et al. (2005) The tortoise and the hare II: relative utility of 21 noncoding chloroplast DNA sequences for phylogenetic analysis. American Journal of Botany, 92(1), 142–166.

Sonet G, Jordaens K, Nagy ZT, et al. (2013) Adhoc: an R package to calculate ad hoc distance thresholds for DNA barcoding identification. ZooKeys(365), 329–329.

Steinke D, Vences M, Salzburger W, Meyer A (2005) TaxI: a software tool for DNA barcoding using distance methods. Philosophical Transactions of the Royal Society B: Biological Sciences, 360(1462), 1975–1980.

Taberlet P, Coissac E, Pompanon F, Brochmann C, Willerslev E (2012) Towards next-generation biodiversity assessment using DNA metabarcoding. Mol Ecol, 21(8), 2045–2050.

Takano A, Fujita H, Kadosaka T, et al. (2014) Construction of a DNA database for ticks collected in Japan: application of molecular identification based on the mitochondrial 16S rDNA gene. Medical Entomology and Zoology, 65(1), 13–21.

The Angiosperm Phylogeny Group (2003) An update of the Angiosperm Phylogeny Group classification for the orders and families of flowering plants: APG II. Botanical Journal of the Linnean Society, 141(4), 399–436.

The Angiosperm Phylogeny Group (2009) An update of the Angiosperm Phylogeny Group classification for the orders and families of flowering plants: APG III. Botanical Journal of the Linnean Society, 161(2), 105–121.

The Angiosperm Phylogeny Group (2016) An update of the Angiosperm Phylogeny Group classification for the orders and families of flowering plants: APG IV. Botanical Journal of the Linnean Society, 181(1), 1–20.

Vences M, Miralles A, Brouillet S, et al. (2021) iTaxoTools 0.1: Kickstarting a specimen-based software toolkit for taxonomists. BioRxiv.

Wagner DB (1992) Nuclear, chloroplast, and mitochondrial DNA polymorphisms as biochemical markers in population genetic analyses of forest trees. New Forests, 6(1–4), 373–390.

Watano Y, Kanai A, Tani N (2004) Genetic structure of hybrid zones between *Pinus pumila* and *P. parviflora* var. *pentaphylla* (Pinaceae) revealed by molecular hybrid index analysis. American Journal of Botany, 91(1), 65–72.

Wilson JJ, Rougerie R, Schonfeld J, et al. (2011) When species matches are unavailable are DNA barcodes correctly assigned to higher taxa? An assessment using sphingid moths. BMC Ecology, 11(1), 18–18.

Wolfe KH, Li W-H, Sharp PM (1987) Rates of nucleotide substitution vary greatly among plant mitochondrial, chloroplast, and nuclear DNAs. Proceedings of the National Academy of Sciences, 84(24), 9054–9058.

Yamamoto S, Masuda R, Sato Y, et al. (2017) Environmental DNA metabarcoding reveals local fish communities in a species-rich coastal sea. Scientific Reports, 7(1), 40368.

Yamamoto S, Minami K, Fukaya K, et al. (2016) Environmental DNA as a ‘snapshot’ of fish distribution: A case study of Japanese jack mackerel in Maizuru Bay, Sea of Japan. PLoS One, 11(3), e0149786.

Yamaura Y, Oka H, Taki H, Ozaki K, Tanaka H (2012) Sustainable management of planted landscapes: lessons from Japan. Biodiversity and Conservation, 21(12), 3107–3129.

Yao H, Song J-Y, Ma X-Y, et al. (2009) Identification of Dendrobium species by a candidate DNA barcode sequence: the chloroplast psbA-trnH intergenic region. Planta medica, 75(06), 667–669.

Yoccoz NG, Bråthen KA, Gielly L, et al. (2012) DNA from soil mirrors plant taxonomic and growth form diversity. Mol Ecol, 21(15), 3647–3655.

Yonekura K, Kajita T (2003) BG Plants: Japanese name–scientific name index (YList). Retrieved from http://ylist.info

